# Measuring Microbial Community-Wide Antibiotic Resistance Propagation via Natural Transformation in the Human Gut Microbiome

**DOI:** 10.1101/2024.11.26.625464

**Authors:** Nadratun N. Chowdhury, Samuel P. Forry, Stephanie L. Servetas, Monique E. Hunter, Jennifer N. Dootz, Joy P. Dunkers, Scott A. Jackson

## Abstract

1)

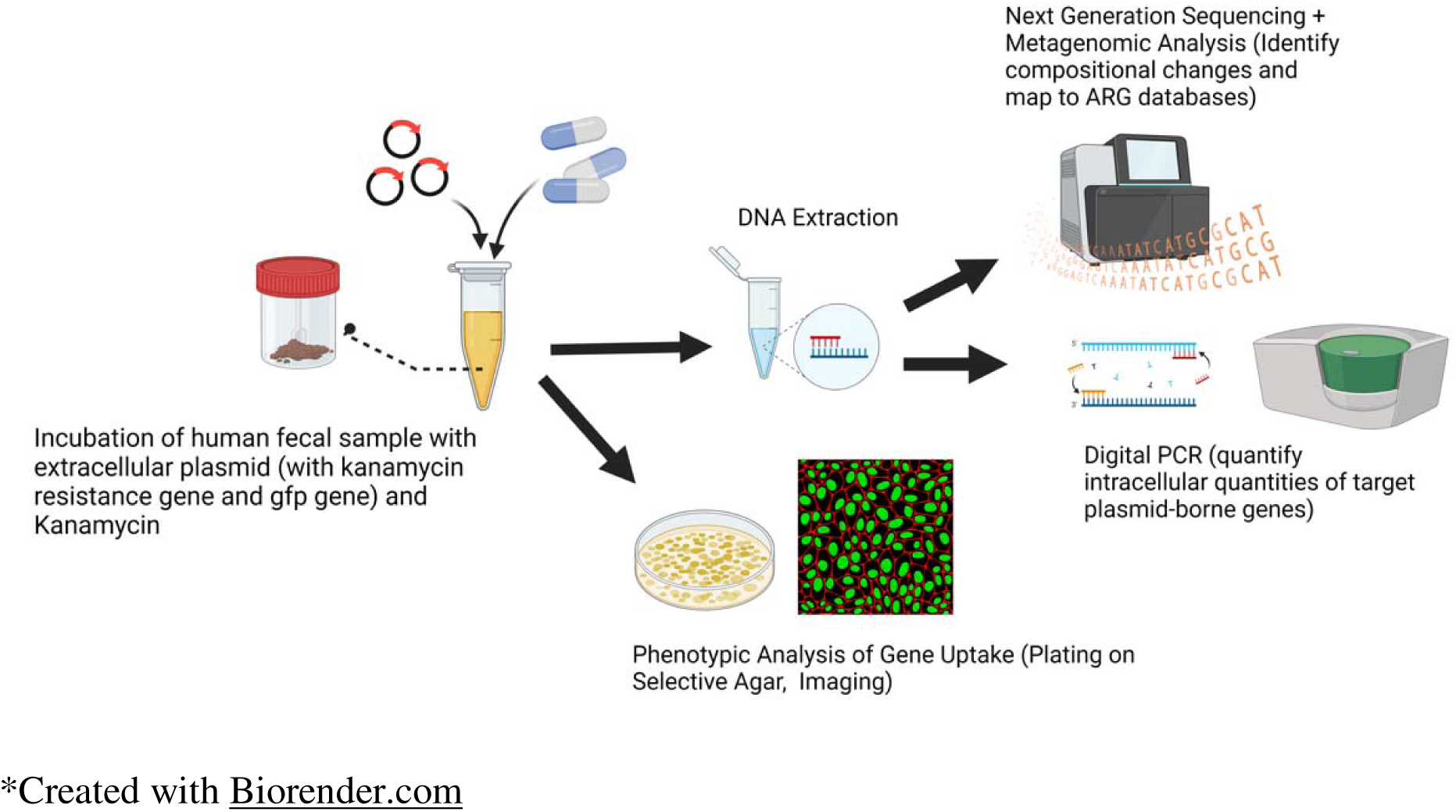

This work explored the role of natural transformation – a mechanism by which bacteria uptake and express extracellular genes - in driving antibiotic resistance propagation in the human gut microbiome. The model extracellular antibiotic resistance gene (eARG) – a plasmid containing a kanamycin resistance (*kanR*) gene and a green fluorescence protein (GFP) gene – was dosed into pooled and homogenized human stool and incubated anaerobically. Cellular uptake of the eARG was assessed via droplet digital PCR, the expression of newly acquired genes was assessed by culturing on selective media and fluorescent microscopy, newly resistant isolates were identified by long-read Nanopore sequencing and the impacts on the taxonomy of the gut microbiome was assessed using shotgun Illumina sequencing. Significant gene uptake of both *kanR* and GFP was quantified in gut microbes, and extent of gene accumulation correlated with background kanamycin levels. Gut microbes dosed with background kanamycin expressed kanamycin resistance acquired by the eARG (as quantified by CFU on kanamycin-containing media). Newly resistant isolates, identified as *Enterococcus faecium* by long-read sequencing, also expressed green fluorescence acquired from the eARG. Though compositional changes of the kanamycin-resistant subpopulation were observed in the gut microbiome in response to eARG and antibiotic exposure, these changes were not reproducible among replicates and trends in taxonomy due to transformation could not be identified. This comprehensive analysis therefore establishes the significant propagation of antibiotic resistance within the human gut microbiome due to eARG exposure, while evaluating the utility of various measurements in characterizing transformation in a complex microbial community.

**Importance:** Infections from antibiotic resistant bacteria in the human gut microbiome are a growing public health concern. Antibiotic resistance may develop in gut microbiota from exposure to environmentally prevalent extracellular antibiotic resistance genes (eARGs). This work explores the impact of eARG exposure on a complex human gut microbial community. It quantifies significant accumulation and expression of eARG-borne genes by endogenous gut microorganisms, thereby demonstrating that natural transformation may play a role in resistance propagation in the human gut. It also demonstrates the highly variable changes in gut taxonomy in response to eARG exposure, implying that eARGs may impact gut composition and therefore downstream human health effects. These data may be useful in characterizing and mitigating resistance propagation in the human gut, and in general, the suite of genotypic and phenotypic measurements used constitute a quantitative framework to characterize the effects of perturbations on complex microbiomes.

## 3) Introduction

The rapid evolution of antimicrobial resistance has led to a major global public health crisis. Infections due to antibiotic-resistant bacteria have contributed to worldwide increases in mortality - estimated at 35,000 deaths/year in the US, 30,000 deaths per year in the EU and even greater numbers in low and middle-income countries in the global south.^1–3^ Such deadly infections may occur in the human gut microbiome - the highly complex and diverse community of bacteria, archaea, viruses, and eukaryotic microbes within the human intestinal tract. The human gut microbiome is host to various opportunistic pathogens, including *Escherichia coli, Klebsiella pneumoniae, Enterococcus faecalis* and *Enterococcus faecium.* Infections from antibiotic-resistant *E. coli*, *E. faecium* and *K. pneumoniae* have been on the rise in last several years.^4,5^ To improve public health outcomes, there is a growing need to prevent or mitigate the propagation of antibiotic-resistant bacteria in the human gut microbiome. To do this, the mechanisms by which resistance spreads in complex microbial communities must be thoroughly evaluated and understood.

The dissemination of widely used antibiotics (for human health and anthropogenic purposes) provides the main pressure for bacterial populations to mutate and develop resistance.^6–8^ However, mobile genetic elements coding for antibiotic resistance – such as plasmids, transposons, integrons, and extracellular chromosomal DNA - may also be agents of spread. Bacteria may acquire mobile genes coding for antibiotic resistance from other bacteria (of the same or different species) through horizontal gene transfer (HGT). HGT may occur via various mechanisms – such as conjugation (in which mobile genetic elements are transferred through a pilus connecting two bacterial cells) and transduction (in which a bacteriophage transfers DNA between bacteria). Perhaps the least studied mechanism, however, is natural transformation - in which bacterial cells uptake extracellular DNA, incorporate this DNA into their genomes and express traits encoded by the DNA. Transformation is an ecologically significant process that has been detected on the bench scale and in situ.^9–13^ Unlike other forms of gene transfer, transformation does not require the presence of a living donor cell, a close genetic relationship between donor and recipient cells, or physical closeness between donor and recipient cells. Therefore, gene networks in a microbial community may be significantly altered based on environmental exposures via transformation.

The transformation process requires bacteria to activate proteins that enable the binding of exogenous DNA to the cell surface, uptake of DNA fragments into the cell, homologous recombination of DNA fragments to the host chromosome (via a recombinase protein called recA), and the translation of exogenous genetic material. Bacteria with this capability are considered naturally competent, and over 80 species of bacteria have been determined on the bench-scale to exhibit natural competence.^23^In most species, the mechanism of competence is inducible, and bacteria may induce competence to repair damaged DNA, obtain nutrients, or incorporate genes that improve their fitness.^10,24–26^ Various environmental factors – such as nutrient limitation or antibiotic concentration – can trigger the activation of competence proteins.^13,14,17^ Some naturally competent species – such as *E. coli* –typically inhabit the human gut.^27^

In addition, the human gut may contain extracellular antibiotic resistance genes (eARGs) released from living and dead resistant bacteria. eARGs are prevalent in the environment and have been quantified in areas that are potential sources for human exposure– including freshwater ^28–30^, sediment ^31, 32^, animal feed operations ^8, 33^, drinking water treatment plants ^8, 34^, and wastewater treatment plants ^35–38^. Though most extracellular DNA quickly degrades when exposed to ubiquitous DNases, detectable levels of dosed eARGs have been isolated and quantified within simulated human gut microbiomes in rats.^39^ This implies that some eARGs may persist long enough in the gut to transform endogenous microbes. Transformed microbes may then disseminate resistance genes to their progeny through vertical gene transfer or to unrelated bacteria via other forms of HGT. Therefore, natural transformation is potentially a significant pathway of resistance acquisition for human gut microbes, but its contribution has not yet been quantified.

Culture-based methods have been previously used to demonstrate that eARGs can genetically transform and confer resistance within bacterial monocultures. ^40–48^ Some studies have quantified transformation in cultured engineered consortia or biofilms of select bacterial strains.^16,49,50^ Culture-based methods, while sensitive, are limited in that they are low-throughput and may only be used to study cultivable strains. In recent years, rapidly developing high throughput culture-independent methods, such as next generation sequencing, have provided the opportunity to collect large amounts of information and comprehensively profile eARGs in environmental microbiomes. These methods are being utilized to study genetic integration of resistance genes in diverse microbial populations– such as in a recent study that evaluated the contribution of eARGs to antibiotic resistance propagation in activated sludge.^43^ In the human gut, conjugative transfer of antibiotic resistance genes has been observed in situ using such culture-independent methods.^51,52^ But there remains a need to make similar community-wide observations of resistance transfer via natural transformation. In this work, the contribution of natural transformation to antibiotic resistance propagation in the human gut microbiome was quantified by dosing a representative eARG into homogenized human stool. Human stool, meant to represent the contents of the gut microbiome, was pooled from multiple donors and verified for homogeneity and stability between samples. The dosed eARG was a synthetic extracellular plasmid deprived of conjugative ability containing a kanamycin resistance gene (*kanR*) and a gene encoding green fluorescence protein (GFP). The microbial community was simultaneously exposed to kanamycin to determine impacts of antibiotic exposure on transformation efficiencies. Genotypic and phenotypic acquisition of kanamycin resistance in gut microbes was assessed using a combination of culture-based and molecular methods. Taxonomic shifts due to antibiotic and eARG exposure was evaluated using next generation sequencing methods, and emergent resistant species were characterized via microscopy and sequencing. This comprehensive analysis informs the current understanding of antibiotic resistance development in the human gut while evaluating the utility of various measurements in characterizing transformation in a complex microbial community.

## 4) Methods

1A) In the first culture step, homogenized human stool is aliquoted into BHI media with and without pBAV1K-T5-GFP (eARG) and kanamycin dosing. Liquid culture is grown anaerobically at 37 °C for 24 hours. 1B) DNA is extracted from stool samples grown in liquid culture. 1C) DNA is sequenced via whole genome sequencing for taxonomic profiling of the treated gut microbiome and analyzed via droplet digital PCR to quantify the dosed eARG.

2A) In the second culture step, liquid cultures grown in step 1 are plated on BHI agar plates, one set on selective plates (with kanamycin) and one set on non-selective plates. Plates are incubated anaerobically at 37 °C for 72 hours. 2B) Agar plates are prepared for molecular analysis by swabbing whole plates into a liquid culture and extracting DNA from this culture, 2C) Samples from the second culture step are analyzed directly from agar plates by counting CFU to determine how treatment with eARG and kanamycin impacted the quantity of resistant cells. Extracted DNA from swabbed whole plates are analyzed via whole genome sequencing to determine the taxonomic profile of the resistant gut microbial population.

3A) Colonies of distinct morphologies are isolated from agar plates containing kanamycin-treated samples from culture step 2, 3B) DNA from isolates are extracted to prepare for molecular analysis. 3C) Isolates, prior to DNA extraction, are imaged via fluorescent microscopy to examine how treatment with eARG impacted expression of green fluorescence. Extracted DNA from isolates are sequenced via long-read sequencing for molecular identification of isolates that emerge after treatment with eARG and kanamycin.

### 4.1) Preparation of materials

#### 4.1.1) Homogenized stool

Human whole stool was obtained from multiple volunteer donors by BioIVT (https://bioivt.com/, Hicksville, NY, USA). All whole stool samples were collected after informed consent under approved IRB protocols at BioIVT. Prior to donation, donors were screened for HIV, Hepatitis B and C, and Syphilis, West Nile virus and *Trypasnosoma cruzi*. Stool samples were collected from multiple volunteer donors on site and immediately stored at -80° C until processing. Upon receipt of all donations, samples were combined with molecular grade water to a final concentration of 100 mg/mL wet weight and homogenized. Samples were then aliquoted in 1mL volumes, shipped to NIST on dry ice, and then stored at -80° C until use.

#### 4.1.2) eARG

The rolling circle plasmid pBAV1K-T5-GFP was used as the model eARG in this study. This plasmid contains genes coding resistance to kanamycin (referred to as *kanR* genes) and a green fluorescent protein (GFP). The *kanR* gene is an aminoglycoside 3’-phosphotransferase (*aph(3’)-IIIa*), which catalyzes the addition of phosphate from ATP to the 3’-hydroxyl group of kanamycin, thereby inactivating antibiotic activity. ^53^ The GFP gene codes for the emission of green fluorescence. The *kanR* gene is used to select for kanamycin resistance and GFP gene is used as an additional marker for phenotypic expression of the plasmid. This plasmid backbone has been previously verified to participate in natural transformation in inter-species studies and shown the ability to positively replicate in a variety of species – such as *Streptococcus pyogens* and *Acinetobacter baumanii*. It also is deprived of its ability to conjugate once inside of bacterial cells. ^50^ Therefore, it was considered an ideal model eARG that may quantifiably transform competent gut microorganisms without confounding effects from conjugation. The plasmid was prepared as described in Text S1 in the Supplementary Information (SI).

#### 4.1.3) Verification of transformability & extractability

*E. coli* was used as a preliminary control to verify the eARG’s transformability. Chemically competent *E. coli* was transformed with pBAV1K-T5-GFP and verified to have the kanamycin resistant phenotype and express green fluorescence, as described in Text S2 and shown in Figure S1 in the SI. The transformed *E. coli* and purified plasmid were then systematically dosed into the stool samples, extracted, and quantified (as described in Text S2 in SI) to verify the ability to adequately extract and measure intracellular plasmid in subsequent transformation experiments. pBAV1K-T5-GFP DNA was successfully recovered from stool samples dosed with transformed *E. coli.* (Figure S2 in SI)

### 4.2) Natural transformation assay

Homogenized stool samples were thawed and 250 µL aliquots were prepared for each sample. Kanamycin sulfate was dissolved into water to make a stock concentration of 5000 g/mL (stored at 4 °C, covered from light). Kanamycin solution was added to stool aliquots, with two aliquots at each of the final kanamycin concentrations – 0, 0.5, 5 and 50 µg/mL. 10 µL of prepped pBAV1K-T5-GFP was added to half of the stool aliquots, such that for each of the four kanamycin concentrations, there was one eARG-free and one eARG-containing sample. This was done to determine the extent to which background antibiotic concentration drove uptake and expression of the eARG. The eARG-free samples with kanamycin were prepared to determine the extent to which the presence of antibiotic caused gut microorganisms to evolve without horizontal gene transfer. The samples for each condition were prepared in triplicate.

Once prepared, the samples were well mixed via vortex. They were then incubated at 37 °C for 24 hours. This incubation period was chosen to be similar to a realistic human gut transit time, which may range between 10 and 73 hours.^54^ Samples were incubated in an anaerobic chamber, which mimics the gut environment. After 24 hours, 1 µL of DNase I was added to each sample to quench transformation, and the samples were incubated another 15 minutes. Samples were then pelleted via centrifugation and resuspended in 100 µL PBS. Intracellular DNA was extracted using the Quick-DNA Fecal/Soil Microbe Miniprep Kit by Zymo Research (Irvine, CA) according to the manufacturer’s instructions, with bead beating on a vortex adaptor at full speed for 40 minutes.

### 4.3) Analysis of gut microbiome upon eARG exposure

#### 4.3.1) Quantification of eARG uptake via droplet digital PCR

Droplet digital PCR (ddPCR) was performed to quantify the GFP, *kanR* and 16S genes in transformed gut microorganisms. The GFP gene was quantified to assess the extent of plasmid presence as an indication of cellular eARG uptake. The *kanR* gene was also quantified to assess eARG uptake but is additionally used to determine the quantity of *kanR* genes present that were not related to eARG exposure. Gene copy numbers of GFP and *kanR* were normalized to copy numbers of 16S, which is a proxy for the abundance of cells present. Because antibiotic treatment likely affects the abundance of cells over time, it was vital to normalize gene copy numbers to 16S gene copies to assess impacts of the eARG or antibiotic. DNA extracted from transformed stool samples was used as template DNA. All experiments were performed on the QX200 Droplet Digital PCR System by Bio-Rad (Hercules, CA). All ddPCR reactions for quantifying GFP and *kanR* genes were performed in 22.5 µL, including the ddPCR Supermix for Probes (No dUTP) by Bio-Rad. The thermal cycling protocol used for these two genes was: 95 °C for 10 min followed by 40 cycles at 94 °C for 30 s and 59 °C for 60 s, followed by 98 °C for 10 min. All ddPCR reactions for quantifying 16S genes were performed in 25 µL, including the QX200™ ddPCR™ EvaGreen Supermix by Bio-Rad. The thermal cycling protocol used for this gene was: 95 °C for 10 min followed by 59 cycles at 94 °C for 30 s and 60 °C for 60 s, followed by 98 °C for 10 min. The sets of forward and reverse primers used are given in the SI (Table S1).

#### 4.3.2) Microbiome profiling by metagenomics (WGS)

Changes in microbial composition in response to eARG and antibiotic exposure was assessed by performing whole genome (shotgun) sequencing. Samples were prepared for sequencing using the NEB Ultra II DNA Library Prep kit for Illumina library preparation (New England Biolabs, Ipswich, Massachusetts). Samples were combined by mass and prepared for sequencing following the MiSeq System Denature and Dilute Libraries Guide (Document # 15039740 v10, Protocol A). Denatured libraries were diluted to a final concentration of 12 pM and combined with 5 % PhiX control (V3 cat# 15017666 from Illumina). Paired-end sequencing was performed on an Illumina MiSeq with 2×150 bp reads (MiSeq Reagent Kit v2 600-cycle, cat #: MS-102-3003). Adapter trimming was done as part of the Illumina MiSeq Generate FASTQ workflow. Fastq files were assembled and taxonomy was classified using Centrifuge WoL: Reference Phylogeny for Microbes Release 1 (available on Github; https://biocore.github.io/wol/) built using 10,575 genomes x 381 genes. ^55^

### 4.4) Post-transformation outgrowths

Transformation assays were also followed up by outgrowths of transformed samples on agar plates containing kanamycin. This was done to determine the quantity of gut microbes that displayed phenotypic expression of kanamycin resistance, and to identify specific strains that may have been transformed. First, homogenized stool samples were prepared with pBAV1K-T5-GFP and kanamycin as described in Section 4.2. For this experiment, samples were prepared only at 0 μg/mL and 50 μg/mL kanamycin, each with and without eARG. Kanamycin concentrations of 0.5 μg/mL and 5 μg/mL were not tested here because 50 μg/mL kanamycin appeared to drive transformation most effectively. Each sample type was prepared in triplicate. All incubations were performed in an anaerobic chamber. After 24 hours, the samples were removed from the incubator and serially diluted. Aliquots of 100 μL of samples from each condition were each spread plated in triplicate on both selective agar plates (Brain Heart Infusion (BHI) agar plates with 2.5% sheep’s blood and 50 μg/mL kanamycin) and non-selective agar plates (BHI-blood plates without kanamycin). Counts on non-selective agar plates provided the baseline count of stool organisms, and counts on selective agar plates provided the count of kanamycin-resistant organisms. Agar plates were incubated in the anaerobic chamber at 37 °C for 72 hours. Ample incubation time was given to allow slow-growing microbes to form visible colonies on the plate.

### 4.4) Analysis of cultured gut microbiome after eARG exposure

#### 4.4.1) CFU count

After 72 hours, plates were removed from the incubator and CFU (colony-forming unit) counts were obtained on both selective and non-selective plates. The fraction of organisms that were kanamycin resistant was determined by taking the ratio of CFU counts on selective agar plates to CFU counts on non-selective plates. Agar plates were visualized on the Scan 4000 by Interscience (Saint-Nom-la-Bretèche, France) and images are presented in SI (Figure S3).

#### 4.4.2) Identifying transformed organisms by metagenomics (WGS)

To determine the taxonomic composition of kanamycin-resistant gut microorganisms, and potentially gain insight into the identities of gut microorganisms subject to natural transformation, colonies from the outgrowths were analyzed by shotgun sequencing. Selective agar plates were swabbed after CFU counts, and cells were inoculated in 500 μL PBS. DNA was extracted from cells using the Quick-DNA Fecal/Soil Microbe Miniprep Kit from Zymo Research (Irvine, CA) and sequenced according to the method described in Section 4.3.2.

#### 4.4.3) Visualization & isolation of colonies

Colonies on selective agar plates were also isolated to identify specific taxa in stool that may have participated in transformation. The outgrowths described in Section 4.4 were performed again, but only on kanamycin-containing selective agar plates. After growth, selective agar plates were inspected for colonies with varying morphologies. Distinct colonies were selected for each sample type, and streak-plated on Blood-BHI agar plates with kanamycin. Streak plates were swabbed into a 50% glycerol solution and stored at -80 °C for further analysis.

#### 4.4.4) Fluorescence microscopy

Isolates from the frozen stock (prepared as described in Section 4.4.3) were inoculated in BHI liquid culture with 50 μg/mL kanamycin and grown overnight. Cultures for each isolate were examined via fluorescence microscopy to determine the morphology of the cells, and whether they expressed green fluorescence as another phenotypic indicator of transformation.

For microscopic imaging, 3 µl sample volumes for each sample were pipetted onto a microscope slide cleaned with KOH and treated with poly-L-lysine. A coverslip was placed on top, tamped down lightly and sealed. The images were collected with a Nikon Eclipse Ti2 E Inverted Microscope equipped with a Hamamatsu (Shizuoka, Japan) ORCA-Flash 4.0 LT+ CMOS camera and a SOLA SE U-nIR solid state light source (Lumecor, Beaverton, OR).

Imaging was performed using a 60X, 1.2 NA plan apochromat water immersion objective with Immersol W (Zeiss, Oberkochen, Germany) as the index matching fluid. Two channel images were acquired using manual focus for each image with these settings: GFP channel (ORCA detector: 2 s integration time, SOLA source: 40%), Differential Inferential Contrast (DIC) channel (ORCA detector: 90 ms integration time, LED source: 10%). The Nikon format files were processed in ImageJ.

#### 4.4.5) Identification of stool isolates

To determine the identity of resistant stool microbes isolated from the transformed community, isolates from the frozen stock (prepared as described in Section 4.4.3) were inoculated in BHI liquid culture with 50 μg/ml kanamycin and grown overnight at 37 °C. DNA was extracted from each culture using the Quick-DNA Fecal/Soil Microbe Miniprep Kit from Zymo Research (Irvine, CA) per manufacturers protocol, and genomic DNA was prepared for nanopore sequencing using the Native Barcoding Expansion 1-12 kit (Oxford Nanopore Technologies, EXP-NBD104). Libraries were sequenced using the Ligation Sequencing kit on the MinION Mk1C (Flow Cell R9.4.1) with the option ‘basecalling’ enabled, with a 48 hour run limit, 1.5 hour pore scan frequency, and 200 bp minimum read length. Sequences were demultiplexed during the run. Barcoded reads were loaded into the EPI2ME Desktop Agent and analyzed using FASTQ WIMP which utilizes Centrifuge version 1.0.3-Beta and a RefSeq database (built 06/08/2021) that includes 56,044 sequences that represent bacteria, archaea, viral, and fungi. FASTQ WIMP was run with the default settings.

### 4.5) Data analysis & visualization

All data points were collected in triplicate. Statistical analysis and data visualization was performed in RStudio. Significant differences between gene copy numbers via ddPCR and CFU counts were determined using R functions for the Kruskal-Walis test (among antibiotic and plasmid treatments) due to the lack of normal distribution. Non-metric multidimensional scaling R functions (Bray-Curtis) were used to compare metagenomic composition (among antibiotic and eARG treatments).

## 5) Results

### 5.1) Accumulation of eARG genes in gut microbial community measured by ddPCR

The abundance of GFP gene copies extracted from transformed cells, normalized to 16S gene copies, significantly increased with plasmid addition in all conditions. A significant increase in GFP abundance in eARG-dosed samples compared to eARG-free samples was observed when no kanamycin was added in the background (p< 0.01*). Increasing levels of background kanamycin, however, impacted the magnitude of GFP gene copy number increase between eARG-dosed and eARG-free samples. The impact of eARG addition on normalized GFP gene copy number was significant at all kanamycin concentrations (p< 0.025* at 0.5 ng/µl Kan; p<0.028* at 5 ng/µl Kan; p<0.006** at 50 ng/µl Kan), and the magnitude of this effect increased with increasing concentrations of background kanamycin. The largest increase was observed when 50 ng/µL of kanamycin was present. Of note is that the population of the gut microbial community, for which 16S gene copies were measured as a proxy, decreased with increasing concentrations of background kanamycin. Therefore, even when accounting for the decrease in the total microbial population by normalizing to 16S gene copy number, the magnitude of GFP gene accumulation correlated with eARG-dosing increased with higher antibiotic doses.

The *kanR* gene was measured at higher levels than GFP in all samples, even those that were not dosed with the eARG. The addition of the eARG led to similar magnitudes of increase in *kanR* gene copies as GFP gene copies, which is to be expected because a copy of each gene is present on the dosed plasmid. Unlike GFP, however, the quantity of *kanR* genes in eARG-free samples increased with increasing concentrations of background antibiotic (p<0.07).

### 5.2) Increase in kanamycin resistant organisms due to eARG exposure

All stool samples transformed in the presence of 0 and 50 ng/µL of kanamycin showed significant growth on agar plates, and the Ratio Resistant CFU (CFU on selective kanamycin-containing agar plates/CFU on non-selective agar plates) is reported for each sample. The Ratio Resistant CFU is a measurement that describes the number of kanamycin-resistant culturable microorganisms as a fraction of the total culturable microorganisms. There was no significant effect of the eARG dosing on the Ratio Resistant CFU when transformation was performed without background kanamycin (p<0.512). Therefore, eARG-dosing did not lead to significant differences in the population of culturable kanamycin-resistant microbes in the absence of the antibiotic.

The overall Ratio Resistant CFU decreased with 50 ng/µL kanamycin treatment as compared to samples with no kanamycin treatment. However, upon treatment with kanamycin, significant differences in the Ratio Resistant CFU between eARG-dosed and eARG-free samples were observed (p<0.049*). The fraction of cells that expressed kanamycin resistance increased nearly 100-fold upon eARG dosing. Therefore, the amount of culturable gut microbes expressing kanamycin resistance, relative to the total gut microbial community, increased upon exposure to the eARG in the presence of background antibiotic.

### 5.3) Identification of strains that may undergo natural transformation in gut microbiome

Outgrowths of transformed stool microorganisms on kanamycin-containing agar plates led to colonies with various appearances (Figure S3). Three visually distinct types of colonies were discernable. Each of the isolates were imaged, and as their colonies appeared distinct, their morphologies did as well (Table 1). In total, three distinct colony morphologies were identified from the outgrowths. Two of these morphologies appeared to be present in both eARG-dosed and eARG-free samples, and one distinct morphology appeared only in eARG-dosed samples. Colonies of similar morphologies between eARG-free and eARG-dosed samples were placed in the same morphology type category.

**Table 1.**
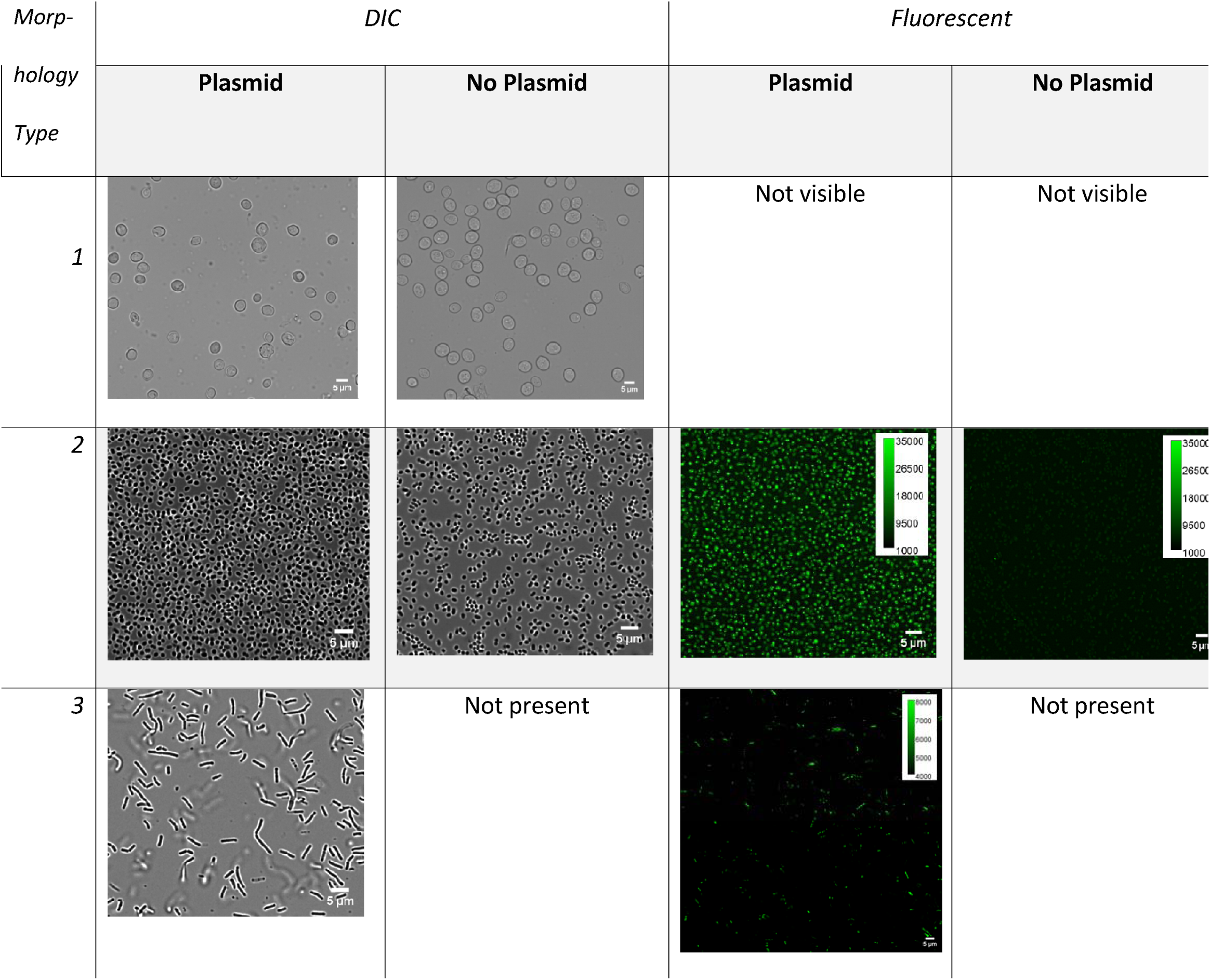
Visualization of bacterial isolates of three distinct morphology types grown on kanamycin-containing agar plates after treatment with kanamycin, with and without pBAV1K-T5-GFP Isolates are visualized on a Differential Inferential Contrast (DIC) channel and a fluorescent channel. Isolates are indicated as “Not visible” when they were present on agar plates but not visible under the microscope, and “Not present” when they were not present on agar plates.

| Morphology Type | DIC |  | Fluorescent |  |
| --- | --- | --- | --- | --- |
|  | Plasmid | No Plasmid | Plasmid | No Plasmid |
| 1 |  |  | Not visible | Not visible |
| 2 |  |  |  |  |
| 3 |  | Not present |  | Not present |

Isolates from colonies of the first morphology type were imaged, and appeared to consist of cells that were circular and large (with a diameter 5-7 µm).These cells most closely identified with a yeast, *Saccharomyces cerevisiae,* with 94.6% and 95.6% of Nanopore sequencing reads matching for the eARG and eARG-free samples, respectively (Table 2). Green fluorescence was not detected in either case, indicating that while this organism expressed resistance to kanamycin, it did not express green fluorescence.

**Table 2.**
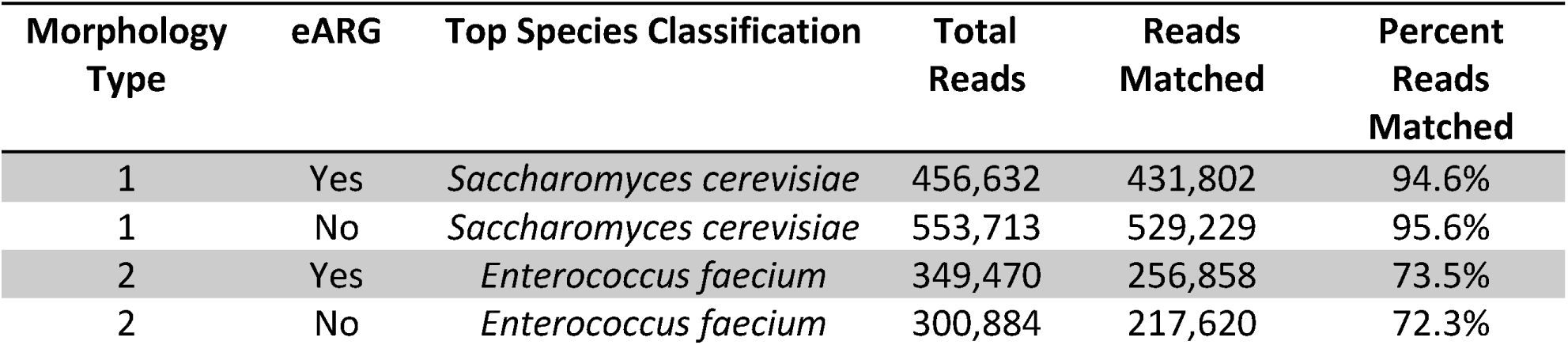
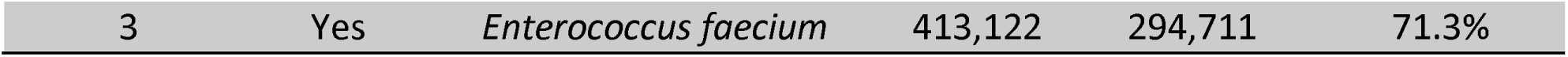
Identification of bacterial isolates grown on kanamycin-containing agar plates after treatment with kanamycin, with and withoutpBAV1K-T5-GFP, by Nanopore sequencing.

Isolates from the second colony morphology type visualized under the microscope appeared to consist of smaller, circular microorganisms with diameters of about 1-2 µm. Morphology type 2 was also found in both eARG-dosed and eARG-free samples. Isolates of this morphology type most closely matched the genus group *Enterococcus/Lactococcus,* with 81.4% and 80.3% of Nanopore sequencing reads matching for eARG-dosed and eARG-free samples, respectively. The closest species match was *Enterococcus faecium,* which had a 73.5% and 72.3% read match for eARG-dosed and eARG-free samples. Green fluorescence was detected for isolates of this strain. However, the isolate from the eARG-dosed sample expressed significantly brighter green fluorescence than the isolate from the eARG-free sample. Though these strains express both kanamycin resistance (as indicated by growth on the selective agar plate) and green fluorescence (as indicated by fluorescence microscopy), the levels of green fluorescence expression appear to increase upon eARG exposure.

The third morphology type was found only on agar plates containing eARG-dosed samples. This isolate was imaged and found to be a rod-like microorganism about 5-10 µm in length. This isolate also most closely matched the genus group *Enterococcus/Lactococcus*, with 79.1% of reads matching, and the species *Enterococcus faecium*, with 71.3% of reads matching. Isolates of morphology type 2 and 3 are possibly different strains of the same genus, which may explain their varied morphology but close genetic match. The expression of bright green fluorescence for the isolate of morphology type 3 was detectable, but by a minority of the microorganisms visualized. Bacteria of this strain, which appear only when treated with the eARG, express resistance to kanamycin, and to a lesser extent, green fluorescence.

When observing the total gut microbial community exposed to kanamycin and the eARG, some reproducible effects on composition were observed with increasing kanamycin addition – such as an increased relative abundances of *Ruminococcus spp, Blautia* spp., and *Eubacterium rectale*, and a decreased relative abundance of *Ligilactobacillus animalis*, *Shigella* spp. and *Enteroccocus faecium* (Figure 4A). Effects of the eARG on the overall gut microbial community composition post-transformation were not discernible via metagenomics. Principal coordinate analysis (PCoA) demonstrated distinct clustering based on kanamycin concentration (Figure 4B). This is to be expected, as antibiotic treatment typically alters the composition of the gut microbiome. Some weak clustering may have occurred based on plasmid treatment, but this effect was marginal. Though changes in gene target accumulation due to plasmid addition were detectable directly after transformation via ddPCR, such changes did not lead to distinct effects on the microbial composition of the stool.

**Figure 1.**
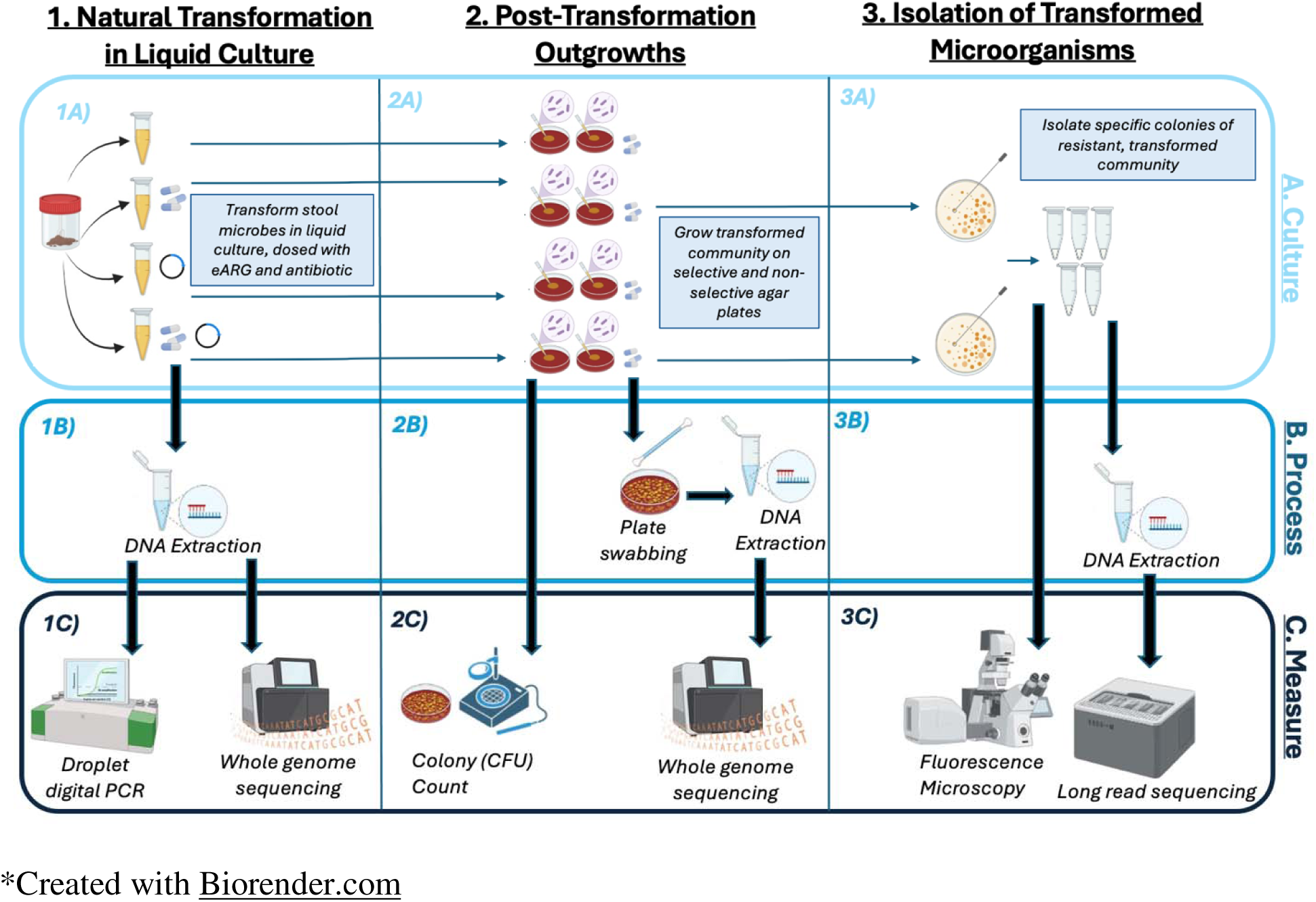
Summary of methods to transform the fecal microbiome with pBAV1K-T5-GFP under kanamycin exposure, and characterize phenotypic and genotypic effects across three culture steps.

**Figure 2.**
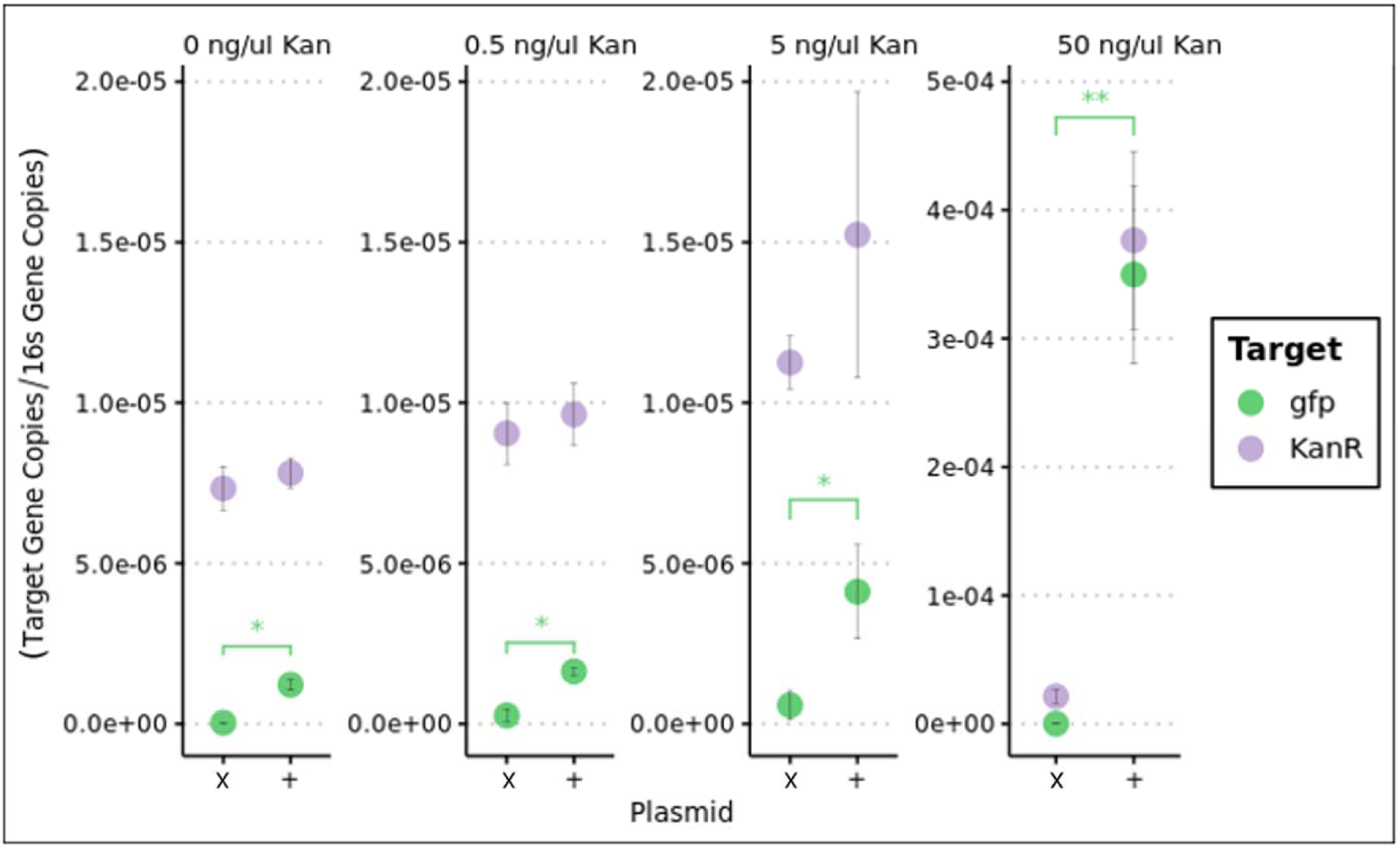
Gene copies of target genes (GFP and *kanR)* from pBAV1K-T5-GFP quantified via droplet digital PCR in DNA extracted from gut microorganisms dosed with pBAV1K-T5-GFP (plasmid) (x = no plasmid dose, + = plasmid dose) and kanamycin, grown anaerobically.

**Figure 3.**
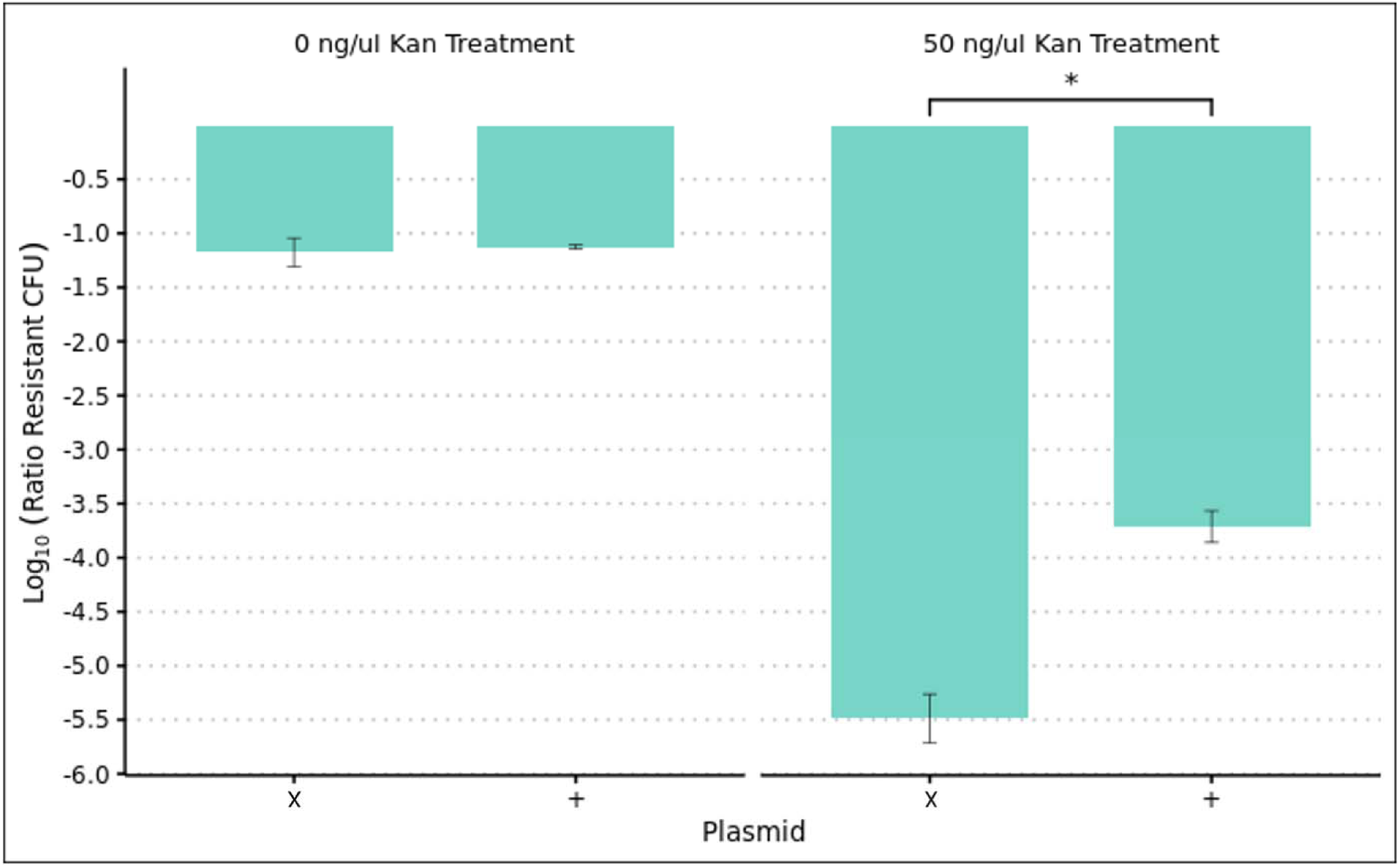
Ratio of CFU on selective kanamycin-containing agar plates to CFU on non-selective agar plates (Ratio Resistant CFU) grown for 72 hours after incubation in liquid culture with pBAV1K-T5-GFP (x = no plasmid dose, + = plasmid dose) and kanamycin.

**Figure 4.**
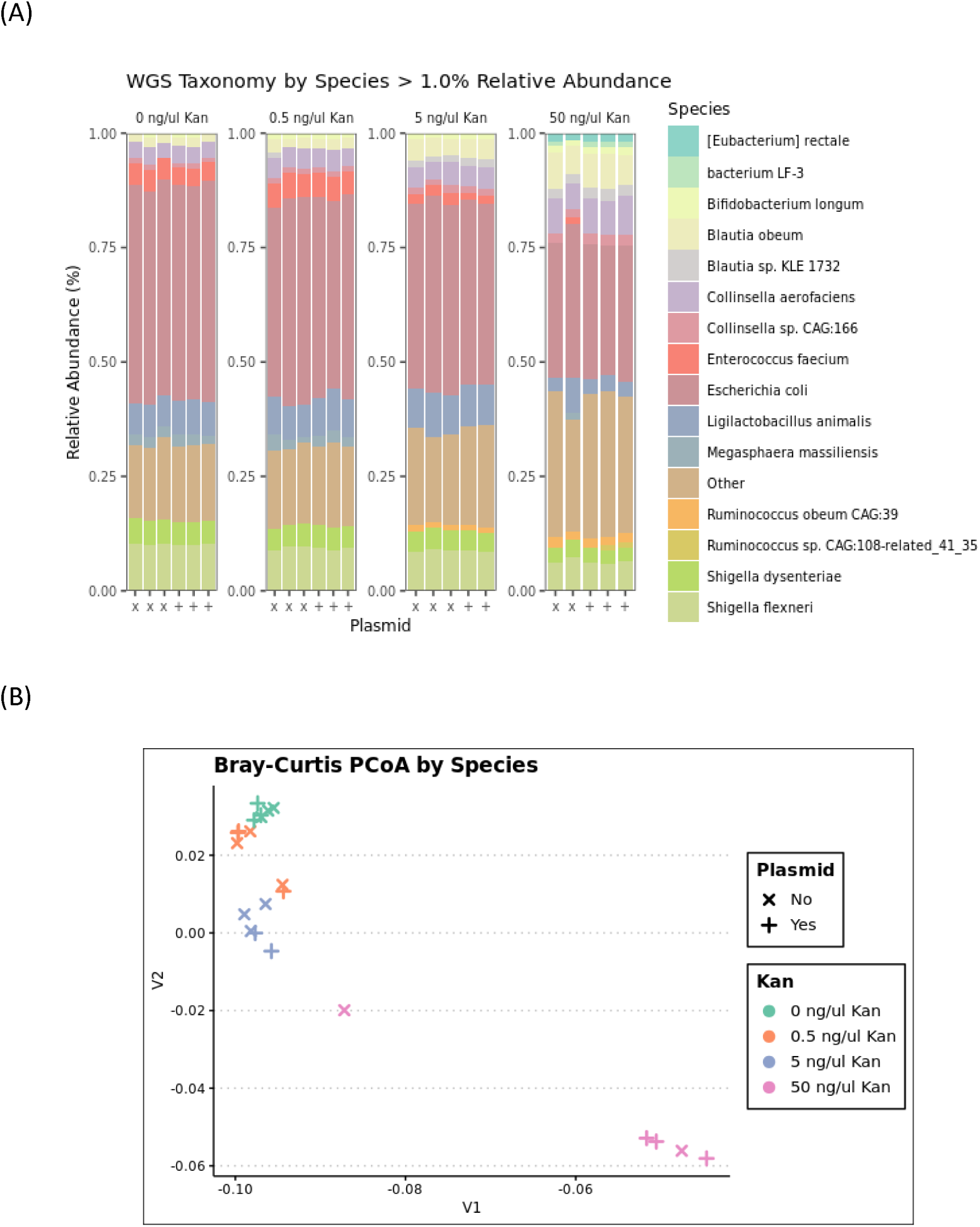
Metagenomic composition changes at a species level in stool population upon 24-hour anaerobic incubation with pBAV1K-T5-GFP (x = no plasmid dose, + = plasmid dose) and kanamycin (Kan) (concentration ng/uL), as shown by A) relative abundance and B) Bray-Curtis principal coordinate analysis (for three replicates).

When observing only the kanamycin-resistant gut microorganisms (swabbed off kanamycin-containing agar plates containing transformed cells), the microbiome did not appear to shift significantly due to eARG addition in samples that were transformed without background kanamycin. (Figure 5A) All samples that were transformed without background kanamycin had similar compositions and clustered together in principle coordinate analysis. (Figure 5B)

**Figure 5.**
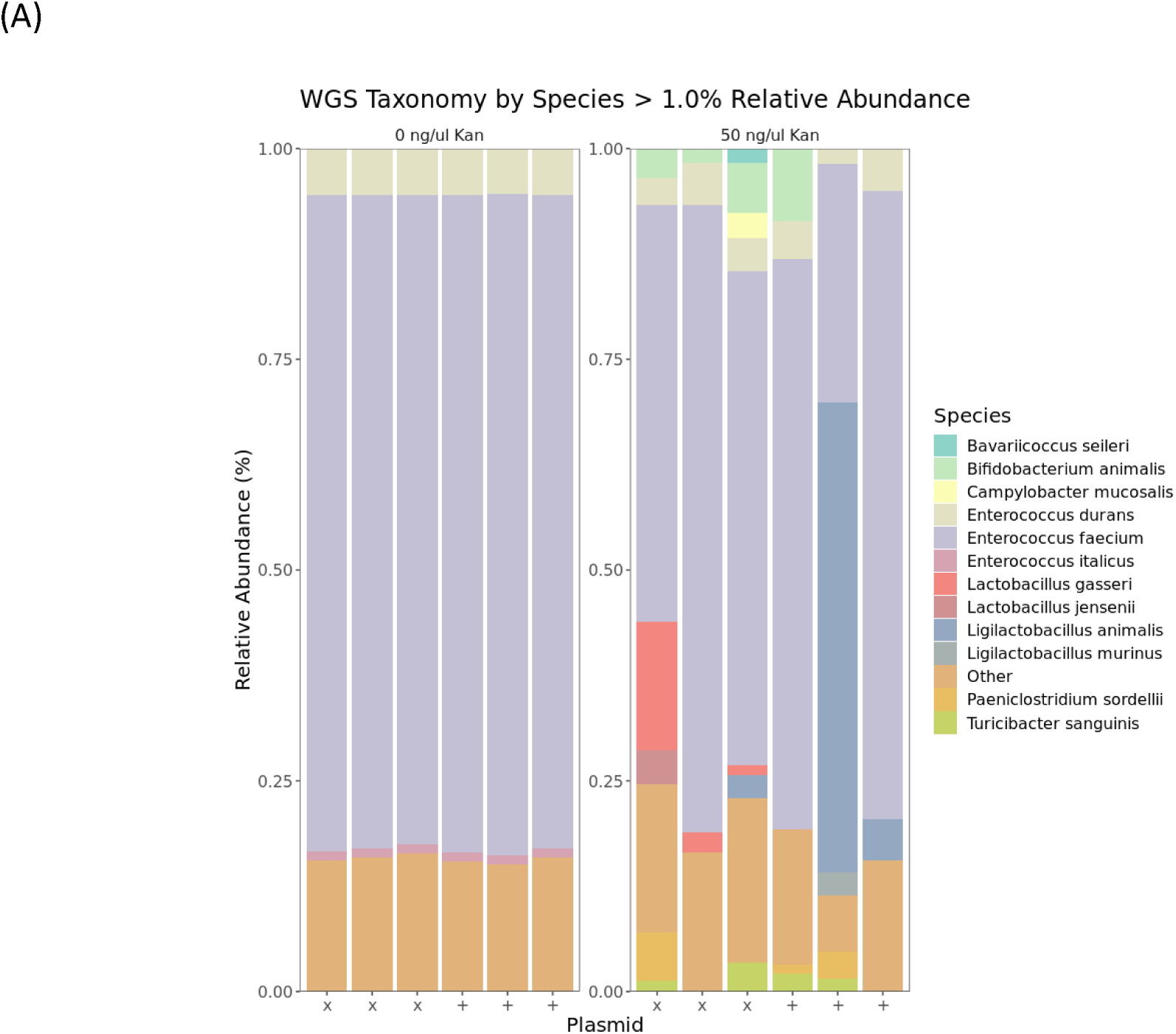
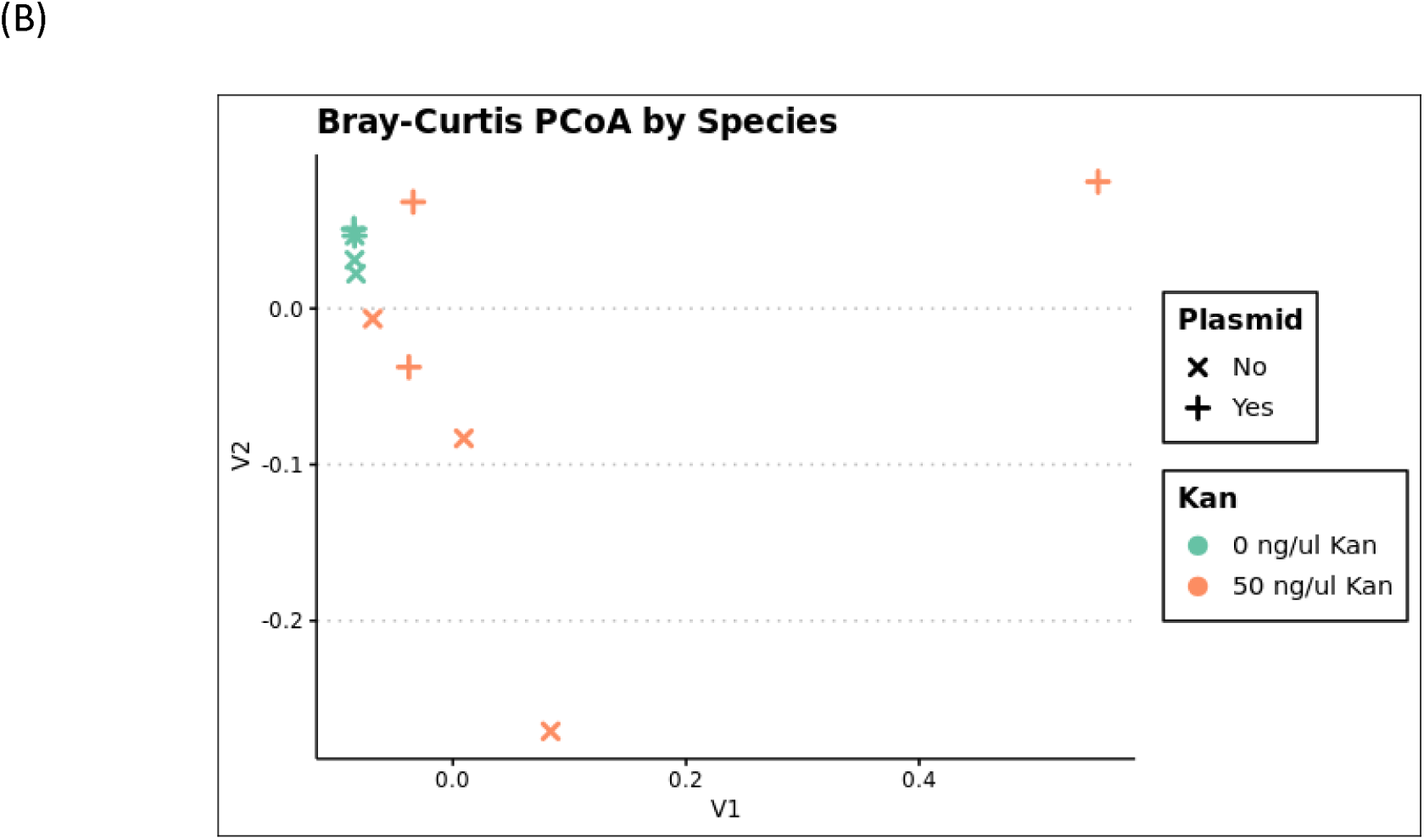
Metagenomic composition changes, at a species level of kanamycin-resistant subpopulation of stool microorganisms (grown anaerobically for 72 hours on kanamycin-containing agar plates) after treatment with pBAV1K-T5-GFP (x = no plasmid dose, + = plasmid dose) and kanamycin (Kan) (concentration ng/uL) (incubated 24 hours anaerobically), as shown by A) relative abundance and B) Bray-Curtis principal coordinate analysis (for three replicates).

The addition of kanamycin at 50 ng/µL led to discernable changes in the taxonomic profile of kanamycin-resistant colonies. As compared to gut microbiome samples not exposed to antibiotic, new taxa were observed in kanamycin treated samples, whether they were dosed with the eARG or not. Such strains may include species like *Lactobacillus gasseiri, Lactobacillus animalis, Paeniclostridium sordellii, Turicibacter sanguinis*, and *Bifidobacterium animalis*, where each showed increases in relative abundance in some kanamycin-treated samples.

Specifically, *Ligilactobacillus animalis* appears in the case of the eARG-treated samples, while *Lactobacillus gasseiri* appears in the case of the eARG-free samples. Regardless, kanamycin-resistant strains do not appear to develop in a reproducible manner, as each replicate demonstrated different compositional makeups (e.g., increases in *Campylobacter* and *Bavilorococcus*). Kanamycin-treated samples did not appear to cluster together in principle coordinate analysis, by eARG-treatment, or overall.

## 6) Discussion

ddPCR results showed significant increases in normalized abundance of GFP and *kanR* gene copies extracted from eARG-dosed transformed cells. Measurable quantities of eARG in the DNA of the gut microbial community imply that gut microorganisms were taking up and accumulating the eARG. The initial uptake of the dosed plasmid likely occurred in cells that were induced into a state of natural competence. This may have occurred in response to environmental factors, such as the limitation of nutrients during the 24-hour incubation period without addition of media.^24, 56^ Once taken up, plasmid-borne genes may have further propagated through the microbial community through other forms of horizontal gene transfer or passed from parent to progeny via vertical gene transfer. GFP and *kanR* accumulation is observed even in samples without background antibiotic, implying that cellular uptake of the extracellular DNA may occur even when the exogenous genes don’t provide a selective advantage.

The extent of GFP and *kanR* accumulation increased with increasing levels of background kanamycin. It is possible that the presence of antibiotic stimulates competence mechanisms in gut microorganisms. Antibiotic exposure has been shown to induce competence in a variety of bacterial species ^57–61^, including *S. pneumoniae* exposed to kanamycin and streptomycin.^62^ An increase in competence amongst gut microbes may have led to an increase of eARG uptake, and the resistance trait may have allowed transformed bacteria to have a competitive advantage in propagation. Further, antibiotics may mediate bacterial SOS responses and increase membrane permeability, which could potentially facilitate extracellular gene uptake.^63–66^

Unlike GFP, the *kanR* gene was measured in significant quantities in samples with no eARG addition, possibly because kanamycin resistance genes are present naturally in gut microbiota, as has been demonstrated in previous studies.^67–70^ Of note is that the *kanR* gene quantity in eARG-free samples increased with increasing concentrations of background antibiotic. This implies that antibiotic exposure drives the cellular acquisition of the *kanR* genes through methods other than uptake of the dosed plasmid. Gut microbes may obtain resistance traits via random mutation or conjugative transfer of existing *kanR* genes, and such microbes would have selective advantage. In the presence of antibiotic, it is possible that a complex microbial community will activate various mechanisms to develop resistance.

While ddPCR results measured the likely cellular uptake of the dosed eARG, the translation and expression of newly acquired genes was confirmed by CFU counts of the kanamycin-resistant portion of the population. Without background antibiotic, there was no significant effect of the eARG dose on the CFU counts of resistant organisms, which implies that cells that took up the eARG did not activate mechanisms to express the acquired genes without selective pressure. However, when background kanamycin was present (at 50 ng/µL), eARG-dosed samples contained significantly higher numbers of resistant organisms compared to ARG-free samples. This implies that in the presence of antibiotic, natural transformation may occur in the human gut microbiome and reproducibly lead to higher amounts of antibiotic-resistant bacteria.

Though the dosed eARG was expressed by certain gut microbes, observing isolates of kanamycin-resistant strains post-transformation revealed that different strains may react to the dosed eARG in distinct ways. First, there is a subset of inherently kanamycin-resistant organisms in the gut microbial community that do not express genes on the dosed eARG. *Saccharomyces cerevisiae* (Table 2) is an example, as this strain grew on kanamycin-containing media in all conditions and did not express green fluorescence when dosed with the eARG. It is likely that this strain was not driven to a state of competence, as *S. cerevisiae* typically requires artificial methods, such as electroporation or addition of a chemical such as lithium or polyethylene glycol, to become competent.^71^ Also, because this strain had intrinsic kanamycin resistance, it may not have expended the metabolic energy needed to retain and express plasmid-borne genes that would not provide any additional selective advantage.

Not all strains with intrinsic kanamycin-resistance seem to behave this way, however. The *Enterococcus/Lactococcus* strain of morphology type 2 (Table 2) included kanamycin-resistant organisms in both eARG-treated and eARG-free samples, but still showed an increased (brighter) expression of green fluorescence in eARG-treated samples. This implies that whilst this strain had inherently kanamycin-resistant members, it still preferred to retain and express the dosed eARG. It is possible that this strain was more easily activated into a state of natural competence that *S. cerevisiae* - *Lactococcus* and *Enterococcus* are the genera with the highest observed numbers of predicted competence genes among lactic acid bacteria ^72^ ^73, 74^ Further, the increased expression of GFP in this isolate also implies that some transformed bacteria may phenotypically express traits obtained from exogenous DNA, even when such traits don’t provide a selective advantage. When the eARG is taken up into the cell, recombined, and translated – it is possible that the GFP gene is translated within the same segment as the *kanR* gene. This may explain the co-occurrence of these traits. This is realistic, as exogenous DNA segments integrated into host chromosomes by recA are on average about 10,000 bp long^75^ and the dosed plasmid is 3,953 bp long.

Observations of the *Enterococcus/Lactococcus* strain of morphology type 3, which grew on kanamycin-containing plates only if treated with eARG, imply that previously non-resistant strains may possibly turn kanamycin-resistant via transformation by the dosed eARG. Only a small fraction of this strain’s kanamycin-resistant population appears to express green fluorescence, which suggests that transformed cells may preferentially express the *kanR* gene on the plasmid as compared to the GFP gene. This strain is an example of a newly resistant strain that emerges in the human gut microbiome in response to eARG exposure. Whilst enterococci have shown the ability to effectively transfer foreign genes via conjugation^74^, it is unknown whether this strain is particularly likely to be transformed compared to other gut microbes.

Overall, according to metagenomic analysis, the addition of the eARG did not lead to significant changes in the composition of the gut microbial community post-transformation (Figure 4). It is possible that newly transformed cells exist in small abundances and are not detectable amongst the millions of members in the complex microbial community. eARG-induced taxonomic shifts may potentially be more prominent after the acquired gene is able to propagate through the community across time and multiple generations, and effects are therefore not detectable after only 24 hours of incubation. Unlike more sensitive methods such as ddPCR or CFU counts, which may theoretically detect as little as 1 gene copy or 1 cell, metagenomics has less utility in detecting changes in a small subset of the microbial community.

Subsequent culturing of the transformed population, on kanamycin-containing agar for 72 hours, led to compositional shifts in the gut microbial community (Figure 5), and particularly led to a significant reduction in microbial diversity. This reduction may be partly due to the fact that many gut taxa are not culturable on blood-BHI agar. However, this is also likely reflective of the fact that many gut taxa, while tolerant to kanamycin exposure in liquid culture throughout the transformation assay (as shown in Figure 4), do not develop phenotypic kanamycin resistance, and therefore do not grow on kanamycin-containing agar. *E. coli,* for example, did not appear to demonstrate phenotypic resistance to kanamycin as it went from being a dominant taxon in the post-transformation liquid culture community (Figure 4) to not being detected in the agar-grown kanamycin resistant community (Figure 5). In contrast, *E. faecium* becomes particularly dominant in the agar-grown kanamycin resistant population, when it was not present in high relative abundances in the post-transformation liquid culture. *E. faecium* is known to have intrinsically high levels of kanamycin-resistance, but some *E. faecium* strains may have obtained the resistant phenotype from the dosed eARG – as is likely the case with isolates of morphology type 3.

Some new taxa in the agar-grown kanamycin-resistant subpopulation appear to have resulted from eARG exposure (i.e., *L. animalis)*, and some from kanamycin exposure (i.e., *L. gasseiri)* (Figure 5). However, the appearance of these taxa was not reproducible amongst the three replicates of the same treatment (with and without eARG dosing) (Figure 5). Therefore, whether these new taxa are the result of eARG exposure or of antibiotic selective pressure is not discernible. It is possible that taxonomic changes in a microbial community exposed to eARG and antibiotic occur stochastically, and the variability in potential outcomes is too high to capture trends within three experimental replicates. This is reasonable, as antibiotic resistance development in response to selection pressure is subject to chance effects of mutation and genetic drift,^76, 77^ and natural transformation relies on inherently stochastic molecular and cellular events – such as contact between the eARG and the cell surface.^78^Additionally, it is possible that longer-term compositional effects of eARG exposure on gut microbial composition are more reproducible, but not found here because the community was evaluated after only 24 hours of incubation. Therefore, though this study identifies various gut taxa (such as *L. animalis* or *E. faecium*) that express phenotypic kanamycin resistance after eARG exposure, reproducible trends in the evolution of gut taxa in response to kanamycin exposure could not be identified within the limited replicates and incubation time allowed.

Regardless of replicate and time limitations, this work develops a methodological framework for comprehensively assessing the effects of perturbations in a complex microbiome using a suite of accessible microbial measurements. A combination of molecular and culture-based methods – including ddPCR, CFU count, microscopy, short-read and long-read metagenomic sequencing - was required to reveal the complex dynamics of the gut microbiome exposed to extracellular elements. Each method provided insights that overcame the limitations of other methods. For example, while some non-cultivable strains were revealed as potential transformants via short-read sequencing (*L. animalis* or *L. gasseiri*), some resistant strains did not appear at high enough relative abundances to be identified by metagenomics and instead were identified as cultured isolates via microscopy and long-read sequencing (*S. cerevisiae*). And in samples where transformation did not lead to community-wide compositional changes as detected via metagenomics, ddPCR was able to identify trends of significant gene uptake. Therefore, the combination of techniques used in this study results in a comprehensive profile of natural transformation in the human gut microbiome, as has not been previously recorded.

One limitation of the method suite used, however, is that the DNA extraction and metagenomic sequencing platforms used resulted in highly fragmented DNA, which did not allow linkage of mobile genetic elements to their host cells. This is why, in this study, identity of transformed gut taxa could only be confirmed in cultured isolates, and only two transformed strains were identified (Morphology Type 2 and 3 in Table 2). However, it is possible that there are several more transformed organisms that are not culturable. Future studies on HGT dynamics within complex microbiomes could benefit from recently developed platforms (such as Hi-C sequencing – a method in which DNA is cross-linked prior to lysis and extraction^79^) that allow matching of host cells to their mobile ARGs. This would further characterize the extent and direction of resistance evolution in complex microbiomes and expand a comprehensive analysis of antibiotic resistance propagation.

## 7) Conclusions

Currently, understanding of complex HGT dynamics in the gut microbiome is limited and only beginning to develop. Within this landscape, this work provides critical measurements of the frequency of resistance acquisition in response to eARG exposure. Accounting for other potential biological factors by using a homogenous and stable material in a highly controlled experimental design - exposure to an eARG, in tandem with antibiotics, is shown to increase the number of organisms that possess both genotypic and phenotypic resistance traits. Quantifying HGT due to eARG exposure is an essential piece of understanding the multi-faceted evolutionary behavior of the human gut microbiome, particularly regarding antibiotic resistance evolution.

Whilst the experimental design included large doses of both eARGs and antibiotics into the gut microbiome to effectively characterize bacterial responses, it is reasonable to assume that similar processes could occur in humans, as both eARGs and antibiotics are environmentally prevalent, and antibiotics are administered regularly to patients. In addition, because ARGs encoded on mobile genetic elements may persist in the microbial community for up to several months of time post-antibiotic regimen^80, 81^, environmental eARG exposure to the gut microbiome could potentially cause downstream effects, such as facilitating the occurrence of gastrointestinal diseases or other health effects that warrant further study.

Because this study demonstrates that transformation can impact traits expressed within a complex microbiome like the human gut, it is also reasonable to predict that resistance may propagate in similar ways in other relevant microbiomes. The impact of eARGs on evolving resistant populations in other clinical (i.e. skin, vaginal) or environmental (i.e. wastewater, indoor spaces in hospitals) microbiomes is also paramount to mitigating the rise of antibiotic resistance. This work provides a framework of quantitative genotypic and phenotypic measurements with a model eARG, that can practically be used to characterize resistance propagation and could be applied to future microbiome transformation studies.

## NIST Disclaimer

Certain equipment, instruments, software, or materials are identified in this paper in order to specify the experimental procedure adequately. Such identification is not intended to imply recommendation or endorsement of any product or service by NIST, nor is it intended to imply that the materials or equipment identified are necessarily the best available for the purpose.

## Ethics approval and Consent to Participate

All work was reviewed and approved by the U. S. National Institute of Standards and Technology (NIST) Research Protections Office. This study (protocol #: MML-2019-0135) was determined to be “not human subjects research” as defined in the Common Rule (45 CFR 46, Subpart A).

## Data availability

Metagenome sequencing, ddPCR, culture, and imaging results and all code used for analysis are available online (https://data.nist.gov/od/id/mds2-3579).

## Acknowledgements

We would like to acknowledge NIST’s microbiology team, mainly Lisa Stabryla, Ishi Keenum, Jason Kralj, Kirsten Parratt, and Sheng-Lin Gibson for consulting, reviewing, and providing technical expertise for this project. This material is based upon work supported by the National Research Council’s Research Associateship Program (Grant 1644868).

## Authors Contributions

N.N.C, S.P.F., and S.L.S. conceptualized the project and designed experiments. N.N.C, M.E.H, J.N.D. and J.P.D. analyzed samples. N.N.C, S.P.F., and S.L.S. analyzed data. N.N.C wrote the manuscript and prepared figures. All authors reviewed and approved the final manuscript.

