## Supplementary Information for "Measuring Microbial Community-Wide Antibiotic Resistance Propagation via Natural Transformation in the Human Gut Microbiome"

**Text S1.** Preparation of plasmid and visualization by gel electrophoresis

The plasmid pBAV1K-T-GFP was received within *E. coli* cells from Addgene (Watertown, MA). The *E. coli* was streak-plated on Luria-Bertani (LB) broth, from which colonies were inoculated and grown overnight at 37 C on LB agar plates. The agar plates were swabbed to collect a cell culture. Plasmids were prepared from this cell culture using the GeneJET Plasmid Miniprep Kit by ThermoFisher (Waltham, MA) via the manufacturer’s instructions. To determine if pBAV1K-T5-GFP was adequately extracted from within *E. coli* cells, prepped plasmid DNA was first amplified via PCR using the GoTaq Green Master Mix, 2X by Promega (Madison, WI).

The PCR mix included: 12.5 µL GoTaq master mix, 2.5 µL each of 10 µM forward and reverse primer, 2.5 µL template DNA, 5 µL water. Primers used were for GFP as noted in Table S1 below. PCR was performed on the GeneAmp PCR System 9700 (Themo Fisher Scientific, Waltham, MA) at the following conditions: initial denaturation at 95°C for 120 seconds, followed by 35 cycles of 95°C for 30 seconds, 55°C for 60 seconds, and 72°C for 60 seconds, with an extension of 72°C for 5 minutes. PCR products were verified on a 1% Agarose gel (See Fig S2, (A) – Lane 3) and a DeNovix Fluorimeter (DeNovex Inc. Wilmington, DE) before being stored at 4 °C.

**Text S2**. Verifying transformability of eARG

To verify the ability of pBAV1K-T5-GFP to transform cells, One Shot™ TOP10 Chemically Competent *E. coli* from ThermoFisher Scientific (Waltham, MA) were transformed using prepped pBAV1K-T5-GFP plasmid according to the manufacturer’s instructions. Growth of colonies on LB agar plates containing 50 µg/mL of kanamycin and microscopic fluorescent images (see Figure S1, taken as described in Section 4.1.3 in the main text) were used to verify transformation.

**Text S3.** Verifying extraction of eARG from stool

In order to verify the ability to extract intracellular and extracellular pBAV1k-T5-GFP from stool material, both transformed *E. coli* (100 µL of overnight culture at ~ 2*10^8^ CFU/mL, and 100 µL of overnight culture diluted 1:10^6^) and the purified plasmid (10 µL of ~30 ng/ µL solution) were dosed into 400 µL aliquots of homogenized stool. The mixtures were mixed via vortex and DNA was extracted using the Quick-DNA Fecal/Soil Microbe Miniprep Kit by Zymo Research (Irvine, CA). Extracted DNA was amplified via PCR using primers for the GFP gene and gel electrophoresis was performed to determine presence of plasmid, as described in Text S1 above. Gel electrophoresis images are given in Figure S2.

| Target Gene | Forward primer (5’🡪3’) | Reverse primer (5’🡪3’) | Probe | Reference |
| --- | --- | --- | --- | --- |
| *16S rRNA* | CGGTGAATACGTTCYCGG | GGWTACCTTGTTACGACTT |  | 1 |
| *GFP* | TCAATGCTTTTCCCGTTATCCG | CCTGTACATAACCTTCGGGC | ACGGTATGACTTTTTCAAGAGTGCCA | This study |
| *Kan^R^*  *(aph(3’)-IIIa)* | GCCGTGGGAAAAGACAAGTT | TGTGGATTGCGAAAACTGGG | TACAGCTCGCGCGGATCTTTAAATG | This study |

**Table S1.** ddPCR and PCR primers, probes, and amplified regions

| (A) | (B) |
| --- | --- |
| 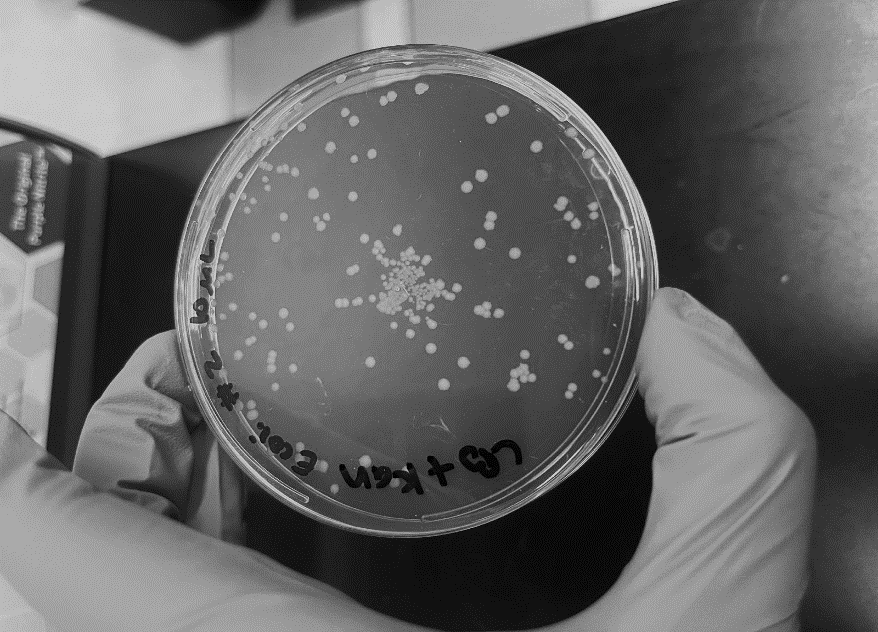 | 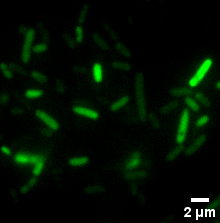 |

**Figure S1.** Chemically competent *E. coli* transformed with pBAV1K-T5-GFP colonies (A) grown on LB agar plates with 50 µg/mL of kanamycin and (B) visualized under a Nikon Eclipse Ti2‑E Inverted Microscope.

| (A) | (B) |
| --- | --- |
| 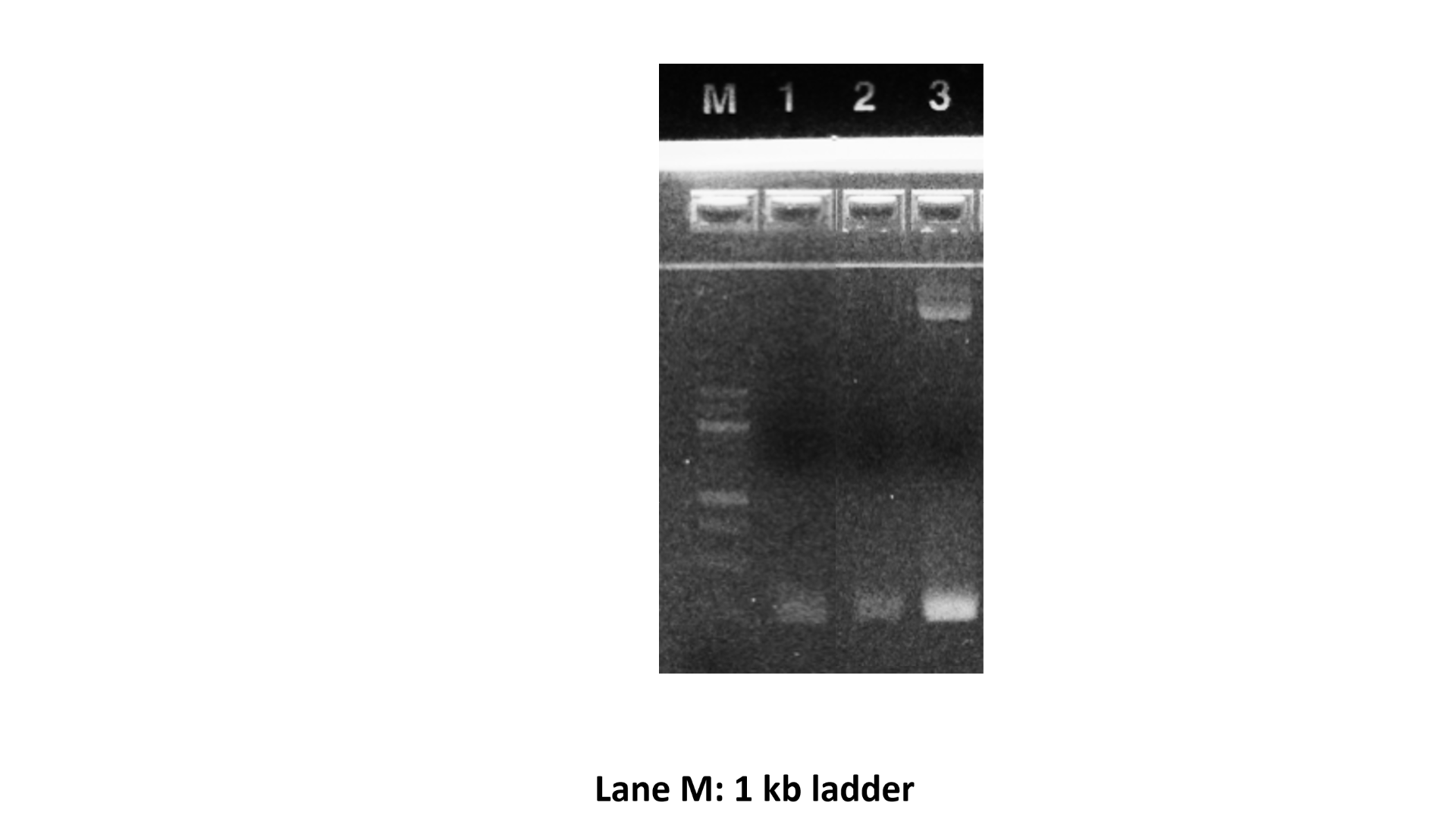 | 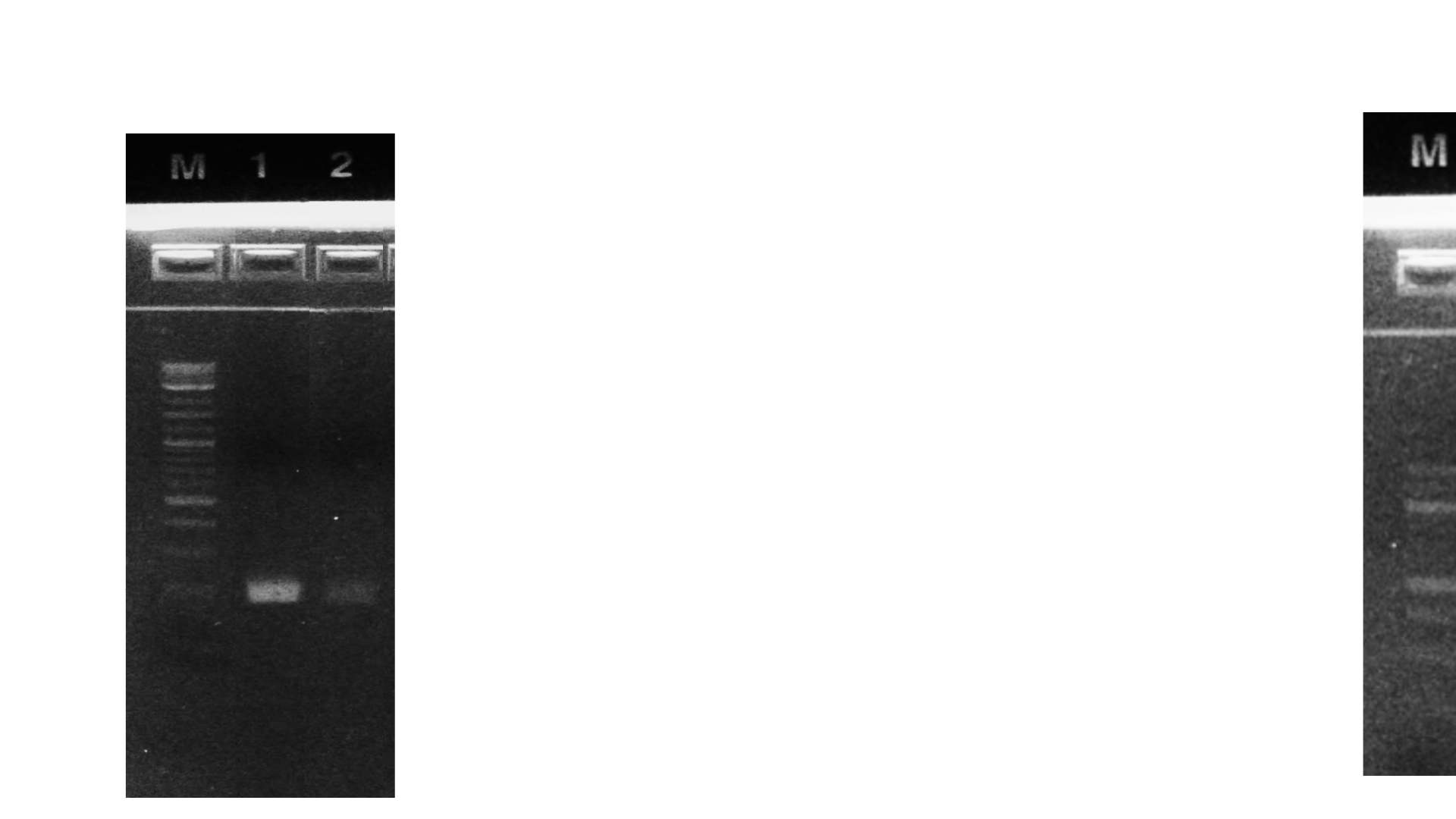 |

**Figure S2.** Verification of pBAV1K-T5-GFP extraction and PCR amplification from *E. coli* and stool.

(A) – Lane M: 1 kb ladder, Lane 1: Stool, Lane 2: Stool with pBAV1K-T5-GFP spike, Lane 3: pBAV1K-T5-GFP grown in *E. coli* (from Addgene).

(B) – Lane M: 1 kb ladder, Lane 1: *E. coli* transformed with pBAV1K-T5-GFP (overnight culture) spiked into Stool, Lane 2: *E. coli* transformed with pBAV1K-T5-GFP (overnight culture diluted 10^-6^) spiked into stool.

| (A) No Plasmid | (B) Plasmid |
| --- | --- |
| 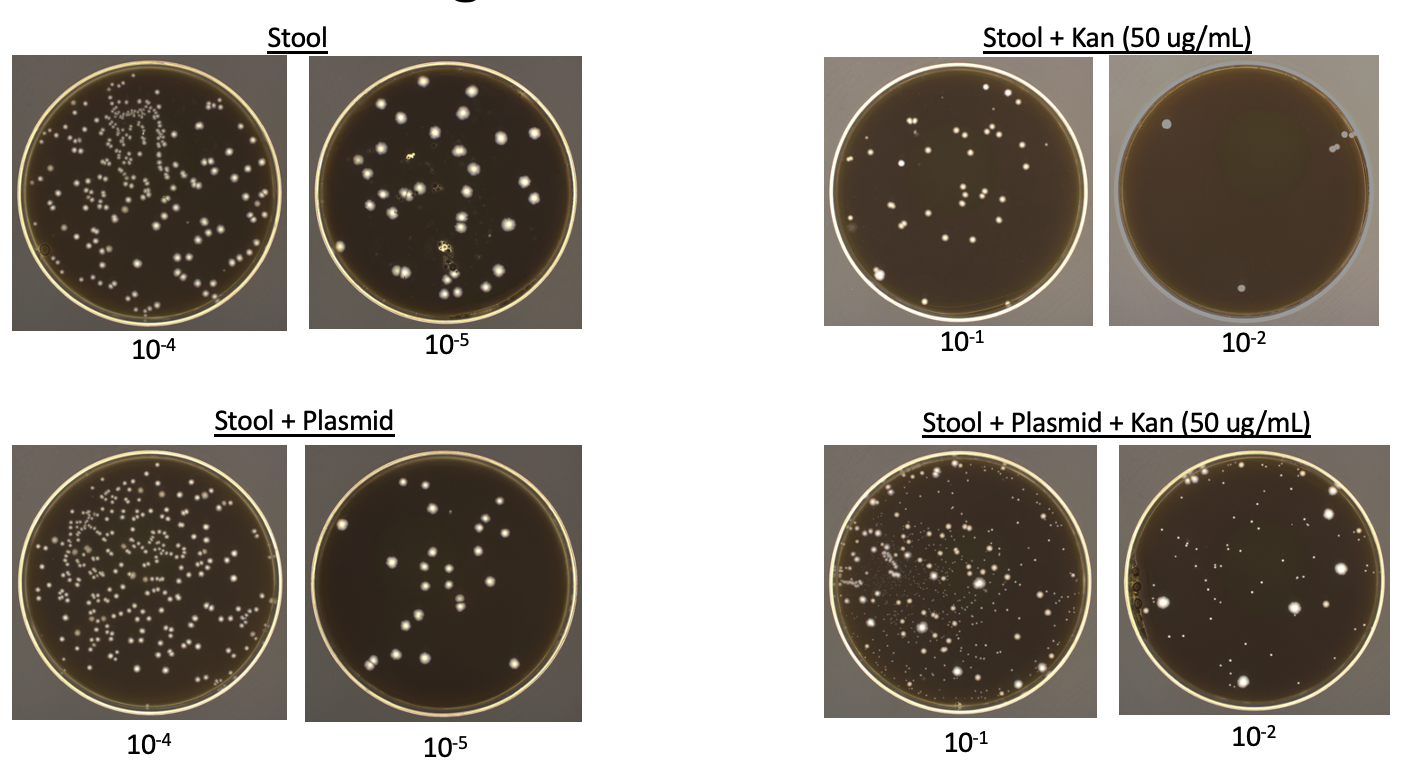 | 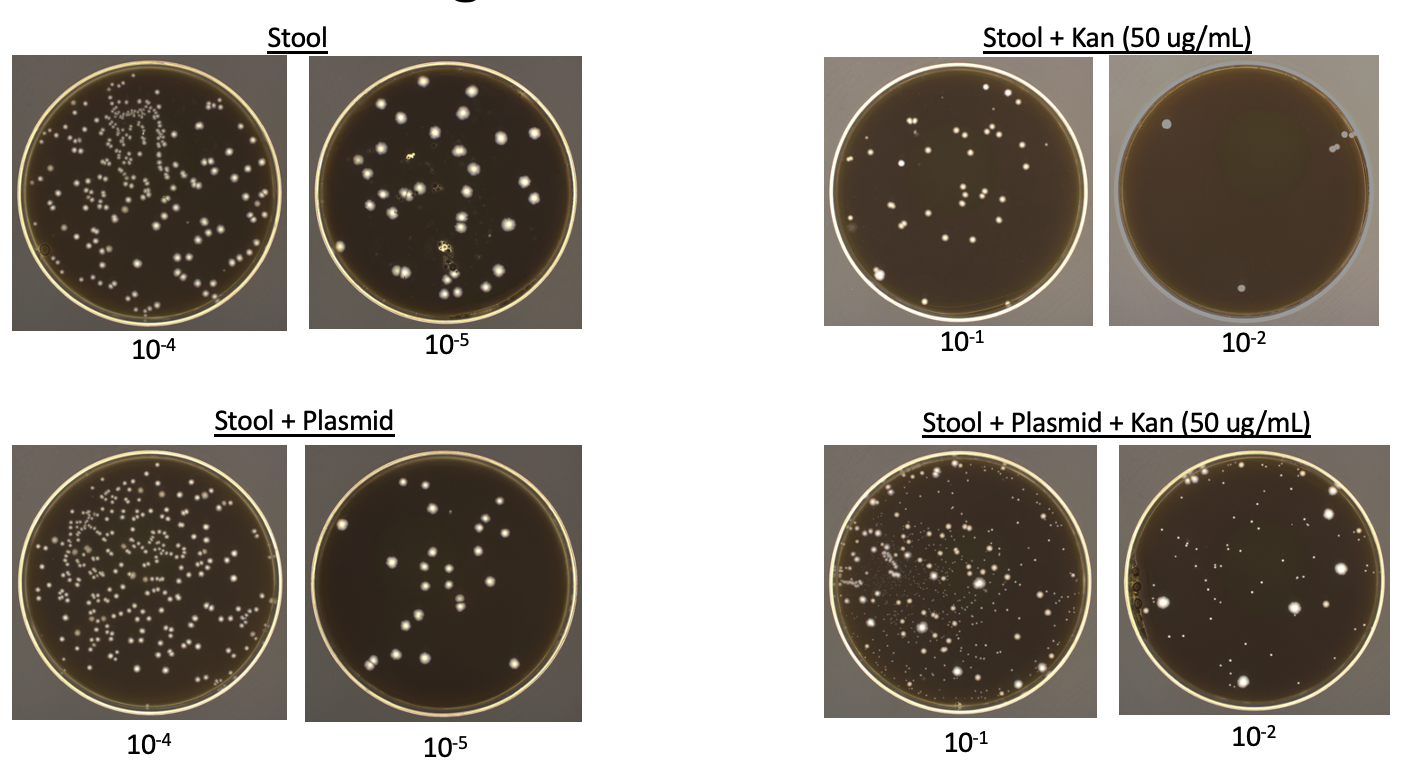 |

**Figure S3.** Change in morphologies of colonies grown with and without plasmid exposure.

(A) - Growth of stool microbes incubated with 50 µg/L background kanamycin on selective agar plates (Blood-BHI with 50 µg/L kanamycin) diluted 1:10^-1^ and 1:10^-2^
(B) - Growth of stool microbes incubated with pBAV1K-T5-GFP and 50 µg/L background kanamycin on selective agar plates (Blood-BHI with 50 µg/L kanamycin) diluted 1:10^-1^ and 1:10^-2^
